# Impairment of homeostatic structural plasticity caused by the autism and schizophrenia-associated 16p11.2 duplication

**DOI:** 10.1101/2025.03.06.641931

**Authors:** MP Forrest, NH Piguel, VA Bagchi, LE Dionisio, S Yoon, M Dos Santos, MS LeDoux, P Penzes

## Abstract

Homeostatic plasticity is essential for information processing and the stability of neuronal circuits, however its relevance to neuropsychiatric disorders remains unclear. The 16p11.2 duplication (BP4-BP5) is a genetic risk factor that strongly predisposes to a range of severe mental illnesses including autism, schizophrenia, intellectual disability, and epilepsy. The duplication consists of a 600 kb region on chromosome 16, including 27 protein-coding genes, with poorly defined effects on neuronal structure and function. Here, we used a mouse model of the 16p11.2 duplication to investigate the impact of this variant on synaptic structure and downstream homeostatic plasticity. We find that 16p11.2 duplication neurons exhibit overly branched dendritic arbors and excessive spine numbers, which host an overabundance of surface AMPA receptor subunit GluA1. Using a homeostatic plasticity paradigm, we show that 16p11.2 duplication neurons fail to undergo synaptic upscaling upon activity deprivation, consistent with disrupted structural plasticity. We also observe that the increased surface abundance of GluA1 occludes further insertion events, a critical mechanism for synaptic plasticity. Finally, we show that genetically correcting the dosage of 16p11.2-encoded *Prrt2* to wild-type levels rescues structural spine phenotypes. Our work suggests that aberrant plasticity could contribute to the etiology of neuropsychiatric disorders.

## Introduction

Neuropsychiatric disorders such as autism spectrum disorder (ASD) and schizophrenia cause life-long impairments to cognitive processing and adaptive behaviors (1, 2). However, the cellular mechanisms that contribute to these symptoms are poorly understood. Large-scale genomic studies have implicated dozens of risk genes in these disorders, many of which are distributed across varied synaptic pathways(3–5). In addition, postmortem studies have highlighted alterations to synapse number and morphology in brain tissue from individuals with neuropsychiatric disorders, implicating these subcellular structures in disease(6). A potential unifying hypothesis in neuropsychiatric disorders is that risk genes commonly impact the process of synaptic plasticity(7, 8), but how this process is affected by defined high-confidence risk variants remains unclear.

The 16p11.2 duplication (BP4-BP5) is a highly penetrant risk variant associated with schizophrenia, ASD, intellectual disability, and epilepsy (9). The duplicated genomic region (∼600 kb) contains ∼27 protein-coding genes, many of which have not been extensively studied in the brain. We have previously shown that correction of proline-rich transmembrane protein 2 (*Prrt2*), a gene encoded in the locus, is sufficient to rescue neuronal network synchrony, seizure susceptibility, and social behavior in 16p11.2 duplication model mice (10). PRRT2 has a multitude of functions at the synapse including regulation of exocytosis via the SNARE complex, tuning neurotransmitter release, and modulating the actin cytoskeleton(11–13). PRRT2 has predominantly been studied at the presynapse, however, it has been identified in proteomic analyses of the postsynaptic density and AMPA receptor complexes (14, 15). Despite a growing understanding of PRRT2 function, the impact of PRRT2 on synaptic phenotypes in the 16p11.2 duplication is unknown.

Pyramidal neurons of the neocortex make thousands of synaptic connections with neighboring cells through small membrane protrusions called dendritic spines(16). Dendritic spines contain all the postsynaptic machinery necessary for neurotransmission, including AMPA receptors, accessory subunits, and their scaffolds(17). Spines are highly dynamic structures that can change shape and size in response to activity-dependent cues, an important aspect of structural synaptic plasticity. Homeostatic plasticity is a long-term form of plasticity, which ensures that global activity levels remain within physiological set points, and is crucial for stable information processing (18). One form of homeostatic plasticity is synaptic upscaling, which occurs during prolonged periods of inactivity. Synaptic upscaling, involves an increase in basal synaptic activity, an accumulation of surface AMPA receptors, and an increase in the size and/or number of dendritic spines(18, 19). Disruption of homeostatic plasticity has been associated with multiple disorders including ASD, intellectual disability, and epilepsy (20–22). Thus, proper brain function appears to require homeostatic mechanisms to effectively tune neuronal and circuit computation.

Here, we assessed the structural modifications to dendrites and spines caused by the 16p11.2 duplication and evaluated the impact of these changes on homeostatic structural plasticity. We find that duplication neurons have excessive dendritic branching and superfluous spine numbers preventing further increases during activity-deprivation. We observe that 16p11.2 neurons have an overabundance of AMPA receptors which limits further insertion events during plasticity. Finally, we demonstrate that the 16p11.2-encoded protein PRRT2, interacts and colocalizes with AMPA receptors in dendritic spines, and correcting its gene dosage restores spine phenotypes in 16p11.2 duplication mice.

Together, our work shows that PRRT2 mediates spine phenotypes in 16p11.2 duplication neurons and that spine overgrowth impairs downstream homeostatic plasticity.

## Results

### Excessive dendritic spine number, size & surface AMPA receptor content in 16p11.2 duplication neurons

Alterations to dendritic spines of cortical pyramidal have been reported in a number of neuropsychiatric disorders, including autism and schizophrenia(6). We therefore set out to determine whether the 16p11.2 duplication (Dup), causes changes to spine phenotypes. We examined the number and morphology of dendritic spines in GFP-transfected pyramidal neurons from mouse cortical cultures (Figure 1A). We found that Dup neurons had a higher density (*p* = 3.4 x 10^-3^) and size (*p* = 1.54 x 10^-2^) of dendritic spines compared to wild-type neurons (Figure 1B-C). We also examined the surface abundance of AMPA receptor subunit GluA1 within dendritic spines, as it is centrally involved in synaptic plasticity and strongly correlates with spine size (23, 24). We first assessed the surface trafficking of GluA1, by overexpressing a GFP-tagged construct in wild type and Dup neurons (Figure 1D). To account for differences in ectopic expression, we measured surface-to-total levels of GFP-GluA1 in soma, dendrites and spines. We found no differences in surface-to-total levels between wild type and Dup neurons in the soma (Figure 1E). However, we observed an increase of surface-to-total levels in dendrites and spines of Dup neurons, indicating a possible alteration to surface trafficking in dendrites (Figure 1F-G). If trafficking of GluA1 is indeed affected in the Dup neurons, we might expect to see changes to endogenous surface levels. Therefore, we live-labeled unpermeabilized neurons with a GluA1 antibody targeting the extracellular N-terminus, as a secondary approach. We found no changes to GluA1 abundance in the soma or dendrites of pyramidal neurons but observed a specific increase of GluA1 in dendritic spines of Dup neurons (Fig. 1H-K). These data reveal that the duplication causes synaptic overgrowth associated with an increase in surface GluA1 levels within spines, indicating a hyperconnectivity phenotype.

**Figure 1.**
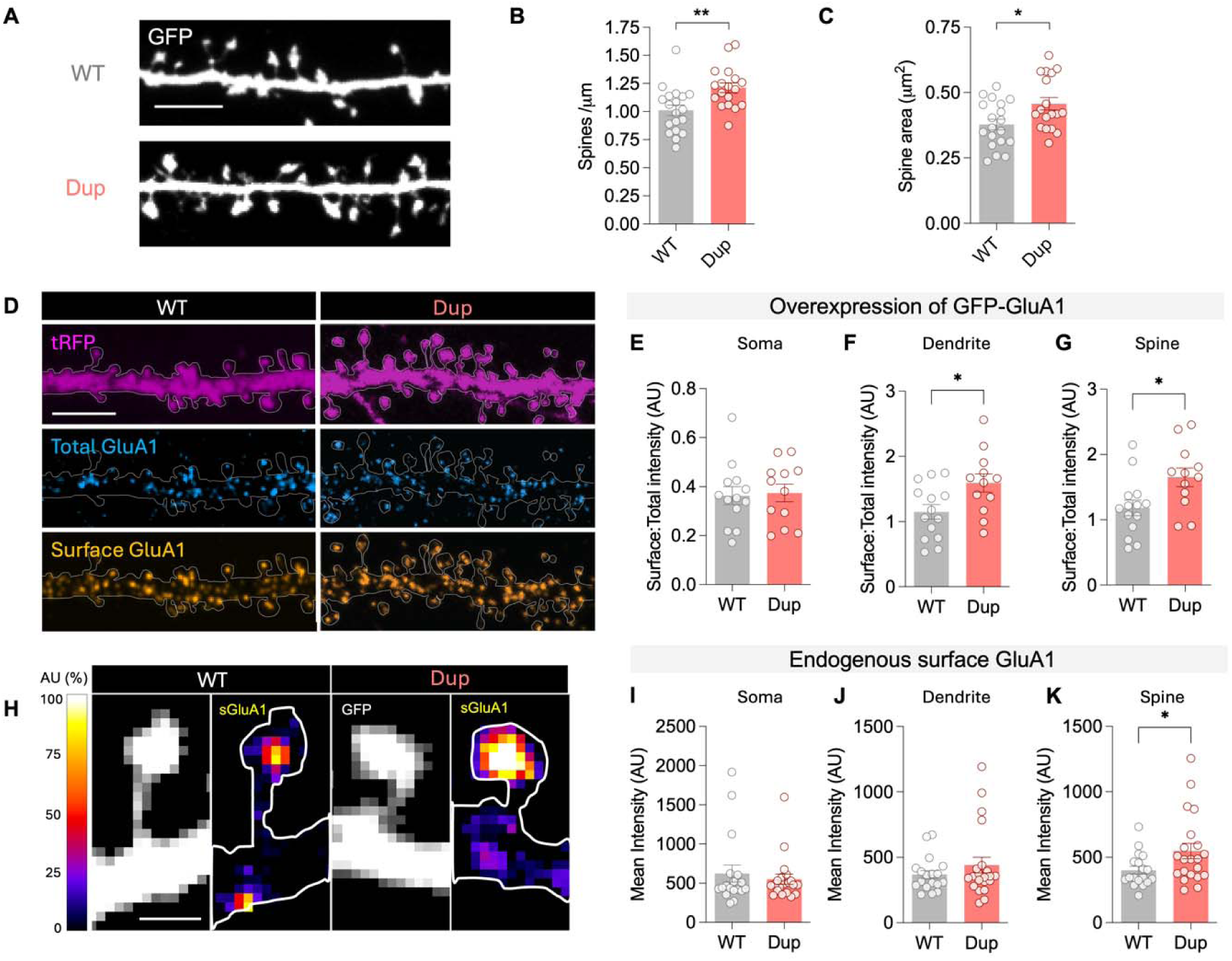
Excessive dendritic spine number, size & surface AMPA receptor content in 16p11.2 duplication neurons. (A) Representative confocal images of GFP-filled dendrites from cultured cortical pyramid neurons. Scale bar 5 μm. (B-C) Increased density and area of spines from cultured pyramidal neurons with the duplication. (D) Representative confocal Images of dendrites with overexpression of a cell fill (tRFP), and total- and surface-labeled GFP-GluA1. Scale bar 5 μm. (E) Surface-to-total ratios of GFP-GluA1 are not altered in the soma (F-G) Surface-to-total ratios of GFP-GluA1 are increased in dendrites and spines of cultured Dup neurons (H) Heatmap of surface GluA1 intensity in dendritic spines. Scale bar 1 μm. (I-K) Surface staining of AMPA receptor subunit GluA1 in cortical neurons shows increased surface levels of GluA1 in dendritic spines of Dup neurons but not in the soma or dendrites. Data are shown as mean ± s.e.m. *p < 0.05, **p< 0.01, two-tailed t-test. Abbreviations: WT, 16p11.2+/+; Dup, 16p11.2dup/+.

### Synaptic hyperconnectivity leads to failure of homeostatic structural plasticity

To determine how an excess of synapses would impact homeostatic adaptation at a structural level, we employed a classical assay for homeostatic plasticity involving a tetrodotoxin (TTX)-based activity deprivation for 48h(25). TTX is a potent sodium channel blocker that inhibits action potential generation and effectively abolishes neuronal activity. As expected, TTX-treatment significantly increased spine size, and nominally increased spine density (Unpaired t-test, p= 0.014; two-way ANOVA, p= 0.0974) in wild type neurons (Figure 2A-C). Strikingly, no changes to spine size or density were observed in Dup neurons, indicating a failure of homeostatic upscaling (Figure 2A-C). Pyramidal neuron dendrites are also susceptible to activity-dependent mechanisms, and we have previously reported that Dup neurons have increased dendritic complexity (26). We therefore wanted to determine the effect of activity deprivation on Dup dendrites. Tracing of GFP-filled dendrites after TTX treatment revealed that wild type dendrites significantly increased in complexity after 48h (Figure 2D-E). However, the already hypertrophic Dup dendrites were not able to further increase their complexity after activity deprivation (Figure 2D-E). Together these data show that Dup neurons are unable to upscale, likely due to a ceiling effect on synaptic and dendritic structure.

**Figure 2.**
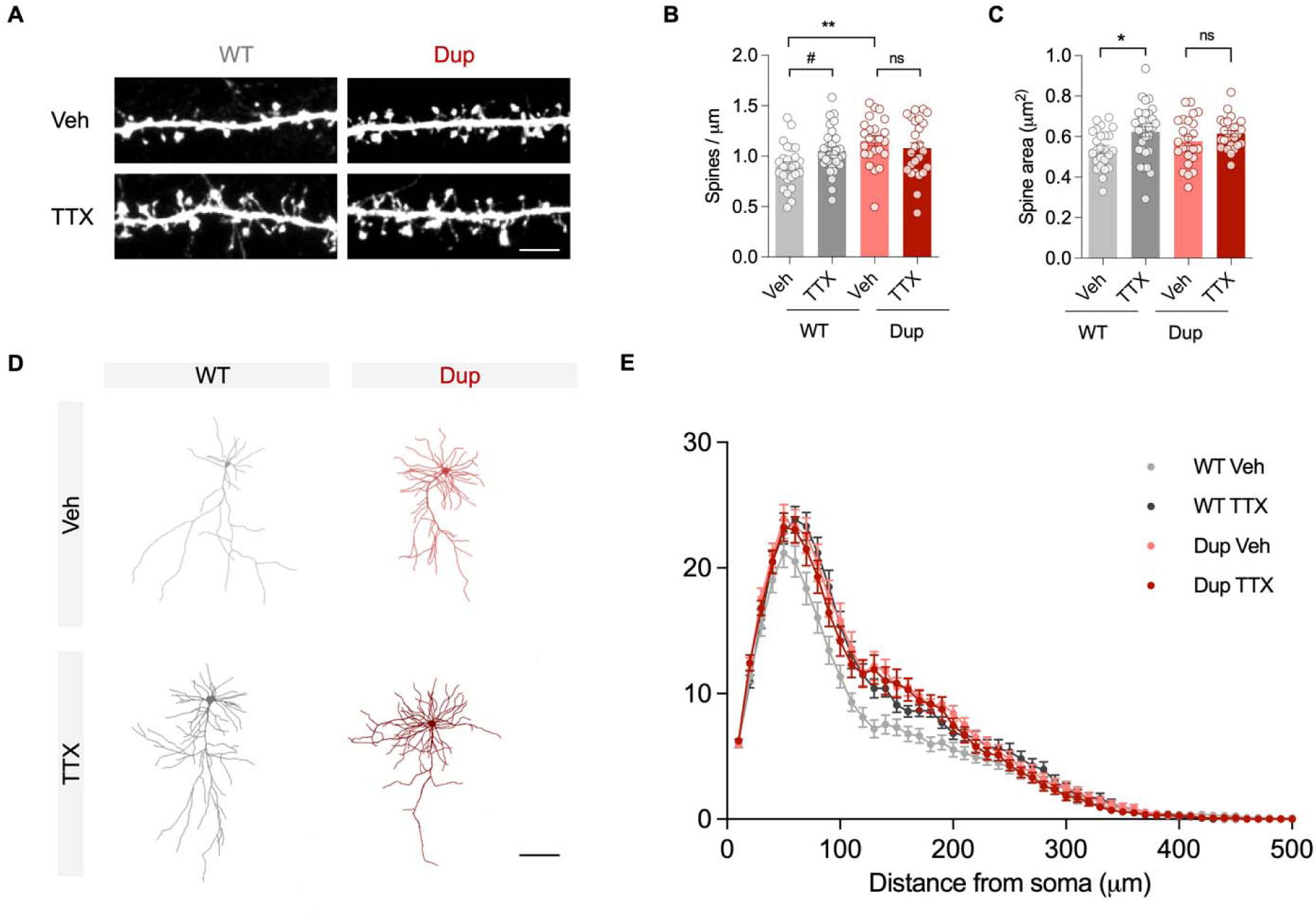
Impairment of homeostatic structural plasticity in 16p11.2 duplication neurons. (A) Confocal images of dendritic spines from cortical neuron cultures treated with TTX (a sodium channel blocker) or vehicle (Veh) for 48h. Scale bar 5 μm. (B) Activity deprivation had a nominally significant effect on spine density in WT but has no effect in Dup neurons (WT Veh vs WT TTX; Two-way ANOVA, p = 0.0974; Unpaired t-test, p = 0.0140). (C) Activity deprivation increases spine size in WT but had no effect in Dup neurons (WT Veh vs WT TTX; Two-way ANOVA with Tukey’s post-hoc test, p = 0.0238). (D) Traces of neuronal dendrites before and after TTX treatment. Scale bar 100 μm. (E) TTX treatment increases dendritic arborization in WT but not in the more branched Dup neurons (Repeated measures two-way ANOVA with Tukeys Post-hoc, Genotype/Treatment x Distance interaction p <0.0001). Data are displayed as mean ± s.e.m. *p < 0.05, **p< 0.01. Abbreviations: WT, 16p11.2^+/+^; Dup, 16p11.2^dup/+^.

### An overabundance of surface AMPA receptors occludes further activity-dependent insertion events

A central mechanism of synaptic scaling is the surface accumulation of postsynaptic AMPA receptors. Given that Dup neurons have increased surface abundance of GluA1 in dendritic spines and there could be a limited number of slots at the synapse(27), we wanted to assess whether Dup neurons were able to recruit new GluA1-containing receptors. Although the exact pathways for surface GluA1 insertion during synaptic scaling are largely unknown, one possibility is that homeostatic plasticity shares surface trafficking machinery with Hebbian plasticity (24, 28). Therefore, we employed an activity-dependent assay to measure AMPA receptor insertion in real-time. To measure new insertion events, we leveraged a superecliptic pHluorin-tagged GluA1 (SEP-GluA1), which is a pH-dependent fluorin that only emits fluorescence at the cell surface when in contact with the extracellular milieu (pH 7), and not does not emit fluorescence in acidified intracellular vesicles (29). We transfected neurons with SEP-GluA1, bleached the entire neuron of surface fluorescence, and measured fluorescence recovery after a brief activity-dependent stimulus (Figure 3A). After 25 min, we found that wild type neurons recovered 15.1% of SEP-GluA1 fluorescence (Figure 3B-C). By contrast, we found that Dup neurons only recovered 8.7% of initial fluorescence, indicating an impairment of GluA1 insertion (p = 0.0153) (Figure 3B-C).

**Figure 3.**
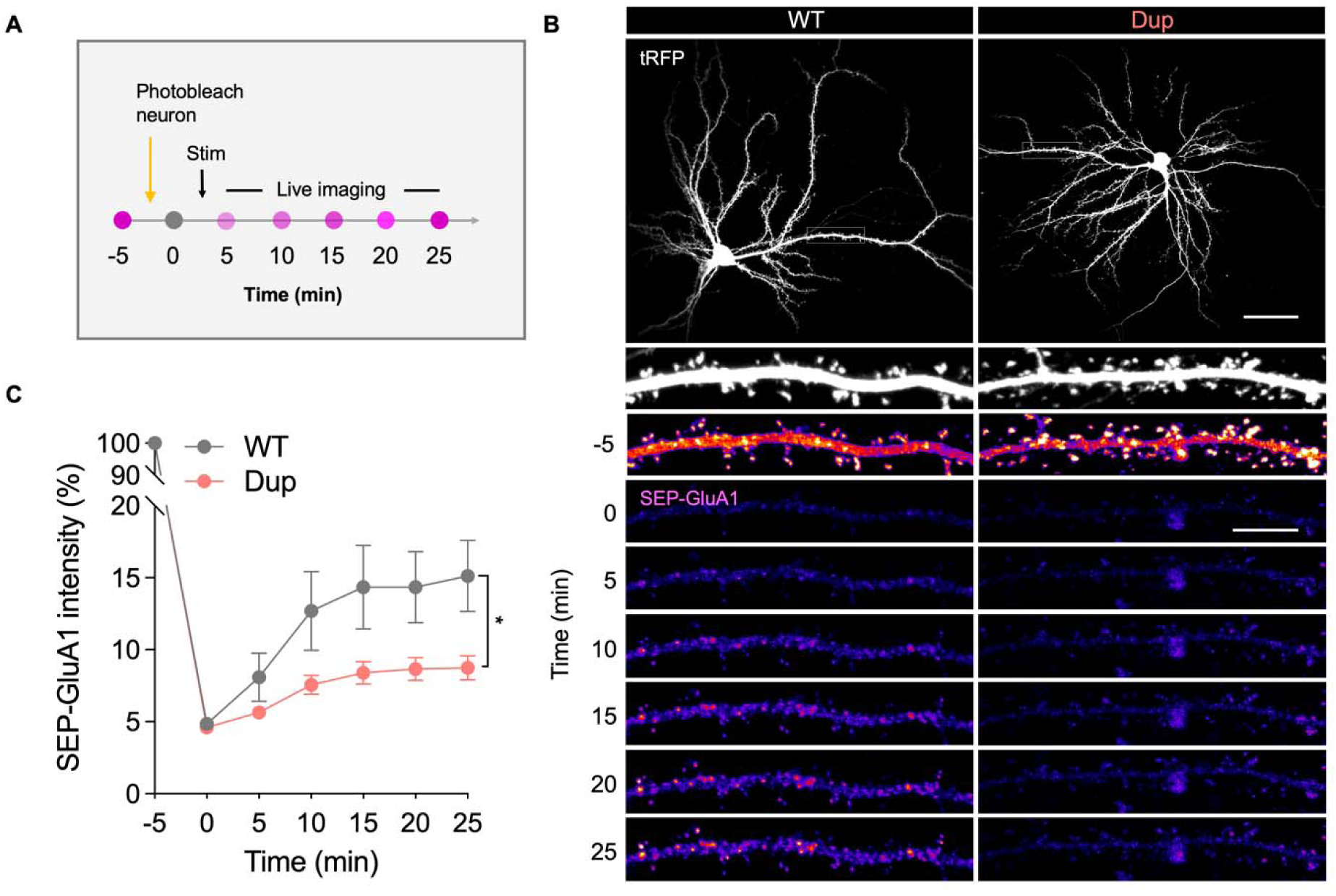
Occlusion of activity-dependent GluA1 surface trafficking in 16p11.2 duplication neurons. (A) Schematic of imaging paradigm used to assess activity-dependent AMPA receptor (GluA1) surface trafficking. Neurons were transfected with SEP-GluA1, extensively photobleached, and fluorescence recovery was measured every 5 min with a confocal microscope. (B) Confocal images of cortical neurons expressing a tRFP cell fill and SEP-GluA1. Bleaching (t=0 min) and fluorescence recovery are shown (t= 5-25min). Scale bar 50 μm (top) and 10 μm (bottom). (C) Fluorescence recovery is reduced in duplication neurons indicating impaired GluA1 insertion (repeated measures two-way ANOVA with Šídák post-hoc test, p = 0.0153). Data are displayed as mean ± s.e.m. *p < 0.05. Abbreviations: WT, 16p11.2^+/+^; Dup, 16p11.2^dup/+^.

### Super-resolution microscopy reveals perisynaptic PRRT2-AMPA complexes in dendritic spines

We hypothesized that AMPA receptor upregulation in Dup neurons could be caused by an increased dosage of PRRT2, a critical regulator of SNARE-mediated exocytosis. Proteomic analyses have shown that PRRT2 is present in mouse PSD fractions and AMPA receptor complexes, suggesting it may have a postsynaptic function (14, 15). However, the precise localization of PRRT2 within postsynaptic compartments and its precise relationship to AMPA receptors and dendritic spines has not been investigated. We used a set of complimentary biochemical and super-resolution imaging approaches, to determine whether PRRT2 has the potential to regulate AMPA receptor trafficking within dendritic spines. First, we confirmed the biochemical association between PRRT2 and AMPA receptor subunit GluA1 using co-immunoprecipitation (co-IP). When overexpressed in HEK293 cells, we found that GluA1 co-immunoprecipitated with FLAG-PRRT2 (Figure 4A). To assess whether this interaction can also occur in native brain tissue, we performed co-IPs from mouse cortical lysates, which revealed that PRRT2 forms a complex with GluA1 *in vivo* (Figure 4B). To further substantiate the association of PRRT2 and AMPA receptors, we searched for proteomic evidence of AMPA receptor subunits and their auxiliary proteins in our previously published PRRT2 interactome dataset. We found a total of 7 out of 17 AMPA receptor-associated proteins were present in our PRRT2 interactome dataset, representing a 2.8-fold enrichment as compared to chance (p=6.45 x 10^-3^) (Figure 4C). Next, we used Stimulated emission depletion (STED) microscopy to assess the nano-localization of PRRT2 and AMPA-receptor subunit GluA1 within dendritic spines, as it has a resolution of 30-70 nm, surpassing conventional confocal microscopy (∼200 nm)(30). Single-plane STED images revealed that PRRT2 and GluA1 co-localize in multiple domains within dendrites and spines of cultured cortical pyramid neurons (Figure 4D). Interestingly, we found that, within dendritic spines, colocalized puncta tended to occur on the periphery of spine heads, suggesting potential sites of GluA1 exocytosis/regulation (Figure 4E). To verify the spine localization of PRRT2-GluA1 nanodomains in the brain, we immunostained Thy1-YFP brain slices and utilized tissue expansion and confocal microscopy to visualize areas of colocalization. We achieved 5-fold expansion factor, thus improving the resolution of confocal microscopy to ∼40 nm. 3D reconstructions of YFP-filled dendritic spines confirmed that PRRT2 colocalizes with GluA1 within these subcellular structures and sites of colocalization tend to occur near spine head membranes (Figure 4F).

**Figure 4.**
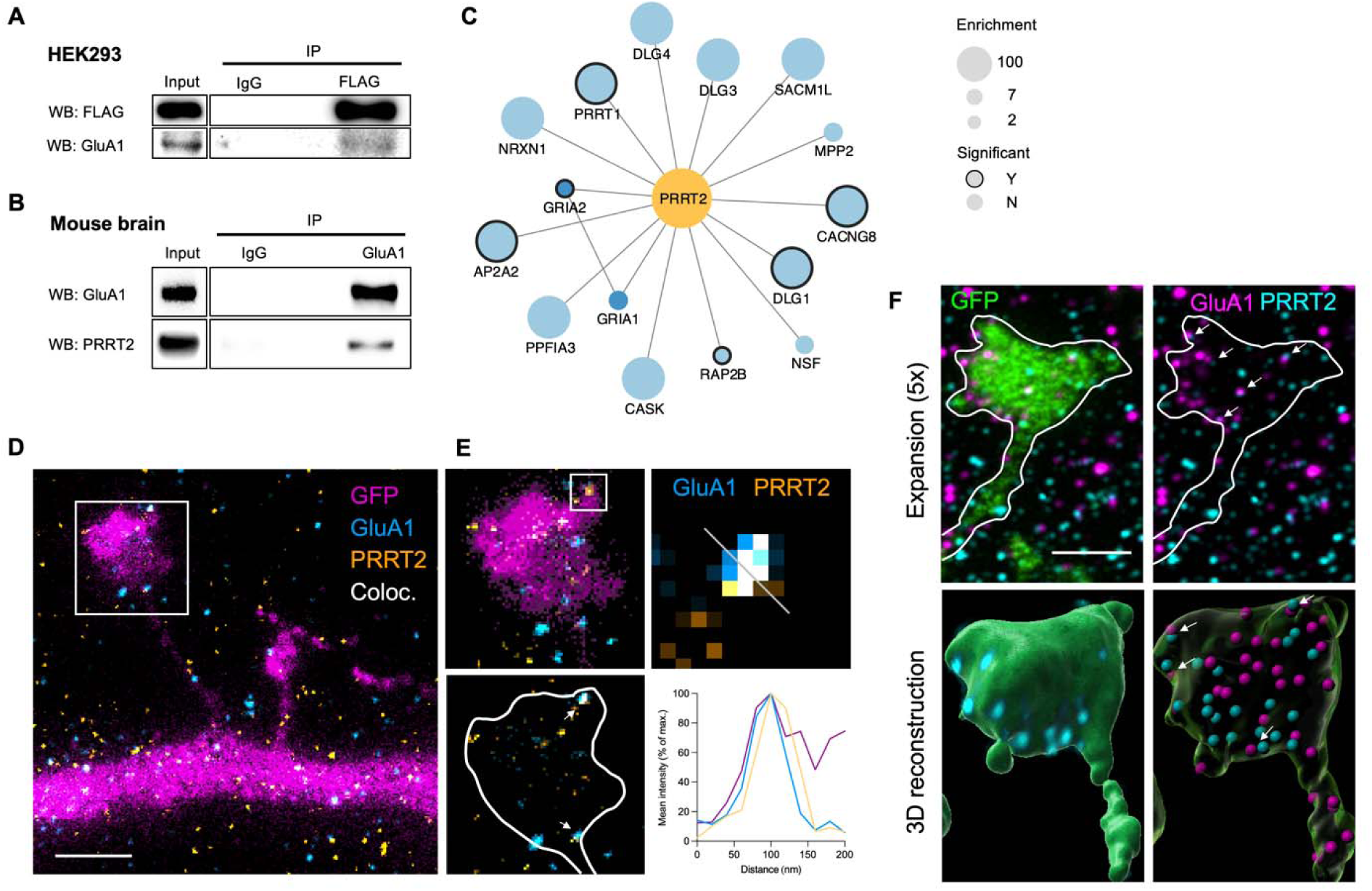
PRRT2 localizes to dendritic spines and interacts with AMPA receptors. (A-B) Immunoprecipitation of GluA1 and PRRT2 in HEK-293 cells (A) and cortical lysates (B) showing that the two proteins co-immunoprecipitate. (C) Protein interaction network showing AMPA receptor complex proteins enriched in a large-scale PRRT2 immunoaffinity purification experiment. This network shows that multiple subunits of the AMPA receptor complex can also co-immunoprecipitate with PRRT2. (D-F) Super-resolution imaging of PRRT2-AMPA receptor nanodomains. (D-E) Single-plane 2D-STED image of GFP-transfected rat neuron showing colocalization of PRRT2 and GluA1 in dendritic spines. Scale bar 1 μm. (F) Expansion microscopy of adult Thy1-YFP mouse brain slice showing colocalization of PRRT2 and GluA1 in dendritic spines of somatosensory cortex neurons. The top panel is a single-plane image and the bottom panel is an Imaris 3D-reconstruction. Scale bar 5 μm.

### Genetic correction of Prrt2 gene dosage rescues synaptic hyperconnectivity in 16p11.2dup/+ mice

We have previously shown that PRRT2 is an important mediator pathophysiology in 16p11.2(10). Nonetheless, the impact of PRRT2 on synaptic and dendritic phenotypes, in the 16p11.2 duplication model has not been determined. To experimentally test the involvement of PRRT2 in synaptic phenotypes, we used mouse genetics to generate a new strain of mice with a full 16p11.2 duplication but normal levels of PRRT2 (referred to here as corrected or “Corr” mice). This was achieved by crossing mice harboring heterozygous *Prrt2* deletions with 16p11.2 duplication mice, as previously described (10). We used cultured cortical neurons transfected with GFP to evaluate spine and dendrite morphology, and immunostained with pre- and post-synaptic markers to assess excitatory synapse density (Figure 5A). This analysis replicated the increased spine density and area in Dup neurons reported in Figure 1 (Figure 5B-C). Strikingly, analysis of neurons from corrected mice revealed that PRRT2 correction fully rescued the aberrant spine phenotypes, implicating PRRT2 as a major regulator of synapse number in Dup mice. (Figure 5A-C). Because dendritic spines host excitatory synapses, we also assessed the density of excitatory synapse markers. Corroborating the spine density analysis, we found that cortical neurons from Dup mice have a higher density of excitatory synapses, measured as the colocalization of the presynaptic marker VGLUT1 and the postsynaptic marker PSD-95 (Figure 5D). In this analysis, correcting PRRT2 expression also reversed the deficits in excitatory synapse density, affecting both the density of pre- and post-synaptic markers (Figure 5D, Figure S1). Interestingly, *Prrt2^+/-^*mice did not have any deficits in spine density or morphology but had a reduced number of VGLUT1 and PSD-95 positive puncta density, similar to *Prrt2* knockout mice (Figure 5D) (31). We also assessed the impact of PRRT2 dosage on previously identified dendritic phenotypes in Dup neurons (26). Although we replicated the increased dendritic complexity in Dup neurons, PRRT2 correction had no effect on dendritic phenotypes, demonstrating its specific effect on dendritic spine and excitatory synapse phenotypes (Figure S2). We next assessed the involvement of *Prrt2* copy number on hyperconnectivity phenotypes *in vivo*, using Golgi-cox staining. We quantified dendritic spines from layer 2/3 pyramidal neurons in the somatosensory cortex, a brain region that presents with a functional hypersynchrony phenotype (10)(Figure 5E). Our analysis revealed an increase in the density of dendritic spines in Dup mice relative to wild type mice, mirroring our findings in cortical neuron cultures (Figure 5F-G). We found no significant differences in spine density between *Prrt2^+/-^* mice and wild type mice. However, correction of *Prrt2* gene dosage restored dendritic spine density to wild-type levels, indicating that *Prrt2* dosage is a critical determinant of synaptic hyperconnectivity in Dup mice (Figure 5F-G).

**Figure 5.**
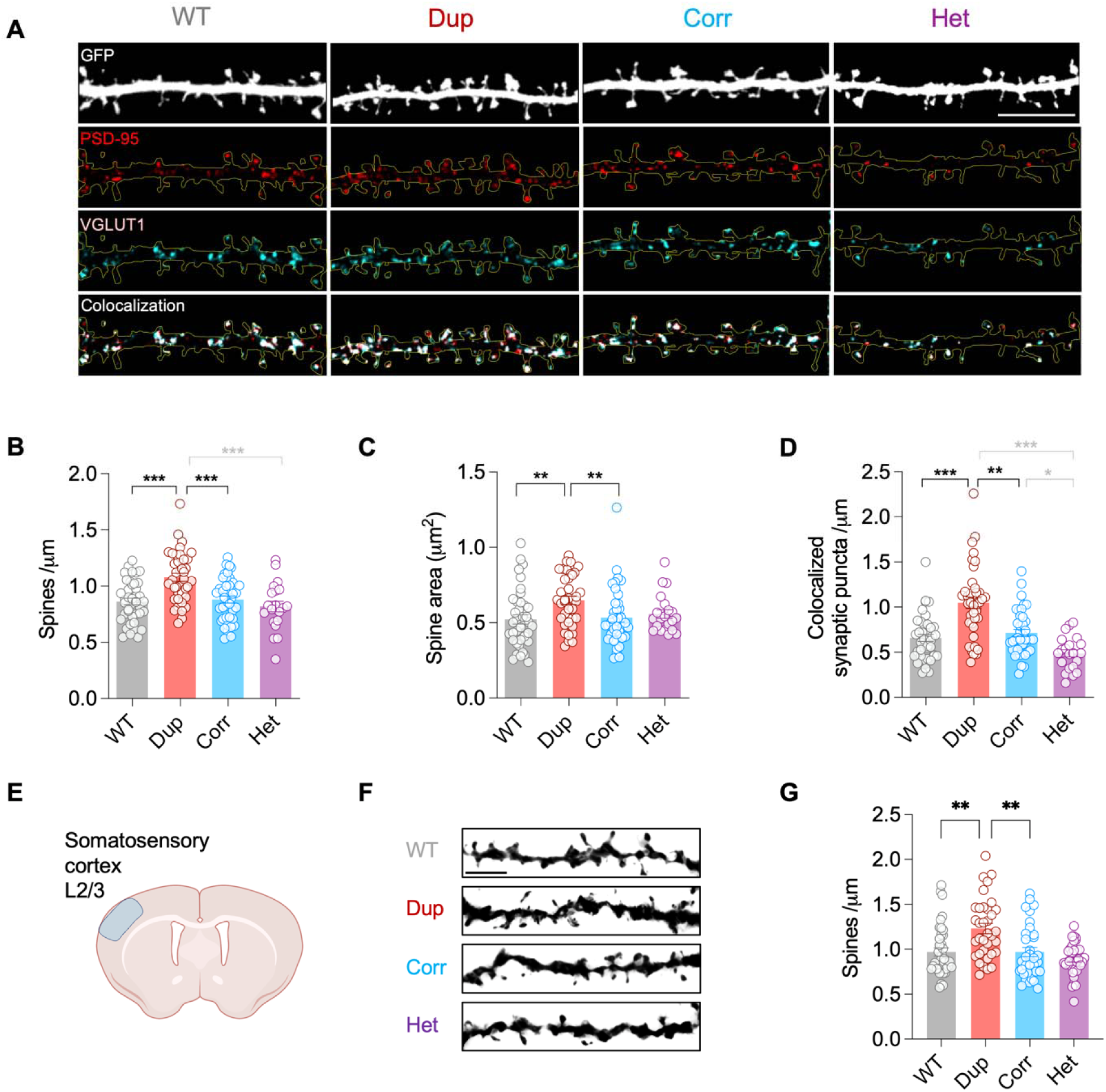
Correction of *Prrt2* gene dosage rescues synaptic hyperconnectivity in 16p11.2^dup/+^ mice. (A) Representative confocal images of GFP-filled dendrites with immunostaining of excitatory presynaptic (VGLUT1) and postsynaptic (PSD-95) terminals. A mask was created around the dendrite to show puncta within the analysis region. Scale bar 10μm. (B) Dup neurons have higher spine density compared to WT, and PRRT2-correction fully restores aberrant spine density. (C) Dup neurons have larger spines compared to WT, and PRRT2-correction fully restores aberrant spine size. (D) Dup neurons have a higher density of co-localized synaptic puncta compared to WT, which is rescued in PRRT2-corrected neurons. (E) Schematic of brain region analyzed for spine analysis. (F) Representative Golgi-Cox stained dendrites. Scale bar 10μm. (G) Dup mice have increased spine density, which is rescued upon PRRT2 correction. Data are displayed as mean ± s.e.m with *p < 0.05, **p< 0.01, ***p < 0.001 (Two-way ANOVA with Tukey’s post-hoc test). Abbreviations: WT = 16p11.2^+/+^, Dup = *16p11.2^dup/+^,* Corr = corrected (*16p11.2^dup/+^*; *Prrt2^+/-^),* Het = *Prrt2^+/-^*.

## Discussion

We have shown novel cellular, synaptic and plasticity-mediated phenotypes associated with a high-confidence risk variant for neuropsychiatric disorders, the 16p11.2 duplication. This work has several implications both at the level of understanding disease mechanisms in the 16p11.2 duplication, and more generally for the role of homeostatic plasticity in neuropsychiatric disorders, which are detailed below.

### Mechanisms of synaptic dysfunction in 16p11.2 duplication mice

Our study revealed that the 16p11.2 duplication causes a structural synaptic hyperconnectivity and failure structural homeostasis. The hyperconnectivity is characterized by an increased complexity of dendritic arbors and an increase in the size and number of dendritic spines. Our experiments support an important role for PRRT2 in the regulation of synaptic phenotypes in duplication neurons. PRRT2 is a synaptic protein tightly associated with synaptic vesicles and SNARE-mediated exocytosis(32). In addition to their role in neurotransmitter release, neuronal SNAREs are critical for the surface trafficking of postsynaptic receptors including GABARs, NMDARs and AMPA receptors, and thus, are also required for plasticity (33–36). Here, we show that PRRT2 interacts with AMPA receptors and forms colocalized nanodomains within dendrites and spines of pyramidal neurons. We also demonstrate that correcting the excessive dosage of PRRT2 normalizes spine density to wild-type levels. One possible explanation for these observations is that PRRT2 regulates the surface expression of GluA1, through the regulation of SNARE-mediated exocytosis, which in turn increases spine size/number. Indeed, previous work has shown that the C-terminal tail of GluA1 interacts with protein complexes required for spine growth/morphogenesis(37). Another hypothesis is that PRRT2 regulates spine numbers by direct effect on the actin cytoskeleton(11). These mechanisms could work in tandem or independently of each other.

In addition to synaptic phenotypes, we have previously shown that the 16p11.2 duplication exhibits hypersynchronous circuitry, increases seizure susceptibility and impaired social behavior(10). Given the strong links between synaptic connectivity and network synchronization (38–40), it is plausible that the synaptic alterations could contribute to network phenotypes in duplication mice, with PRRT2 playing a critical role. Interestingly, a study on 16p11.2 deletion mice found a reduction of spine density in the prefrontal cortex of mice, which was associated with diminished prefrontal connectivity in resting-state functional magnetic resonance imaging (41). These findings suggest that the 16p11.2 locus could regulate synaptic connectivity in a dose-dependent manner.

### Homeostatic plasticity in neuropsychiatric disorders

A potential mechanism contributing to neuropsychiatric disorders is aberrant homeostatic plasticity (20–22, 42). Many genes regulating the structure and remodeling of synapses are implicated in neuropsychiatric disorders, making synaptic plasticity an attractive unifying mechanism that is distributed across risk gene networks. A growing body of work now supports the idea that homeostatic plasticity is affected in genetic models of neurodevelopmental disorders including studies in mouse, rat and human stem cell-derived model systems. Because homeostatic plasticity can manifest in a variety of ways, studies have measured the impact of variants on different types of neuronal outputs including, synaptic currents and intrinsic excitability but very few have investigated structural aspects of plasticity.

Consistent with the idea that homeostatic plasticity mechanisms are distributed across risk gene networks, many different types of risk genes have been implicated including transcription factors, chromatin remodelers, RNA regulatory proteins, and synaptic scaffolds. However, the precise ways in which plasticity is affected in each model, and the underlying pathways involved differ. In *FMR1* knockout neurons, there is no initial difference in basal synaptic transmission however, TTX-based synaptic scaling is abrogated (43, 44). The proposed mechanism for this dysfunction is the lack of synthesis of new GluA1 receptors via FMRP. In haploinsufficient *Chd8* neurons, there is a basal reduction in the frequency and amplitude of miniature excitatory postsynaptic currents, and TTX-based activity deprivation is abolished(45). Finally, in human neurons haploinsufficient for *TCF4*, a model of Pitt-Hopkins syndrome, neuronal activity is reduced and so is the extent of upscaling after TTX treatment(28).

Similar to the phenotypes we describe in 16p11.2 mouse neurons, stem cell-derived neurons with 15q11-13 duplications, another duplication associated with ASD and SZ, manifest with an increased number of dendritic protrusions and excitatory synaptic currents, which in turn impair homeostatic upscaling(46). These findings suggest that synaptic and dendritic overgrowth could structurally prevent homeostatic upscaling, preventing proper synaptic plasticity. Another non-mutually exclusive mechanism is that CNVs in general alter synaptic homeostasis through imbalance of gene dosage, as has been previously suggested(21).

Deficits in homeostatic plasticity could be expected to have several downstream effects on circuit function including aberrant excitation(E)-inhibitory(I) balance, and stability. Interestingly, while chronic inactivity globally upscales synaptic inputs, it is thought to preserve the relative strength of synapses, thereby maintaining previously learned information. It stands to reason that altered homeostatic plasticity could also lead to improper Hebbian learning, which relies on changes to homeostatic set points to continuously encode information. Deficits in homeostatic plasticity could therefore underlie multiple symptom types associated with neuropsychiatric disorders including impairment of cognitive processing, behavioral adaption, and seizures. However, the downstream implications of altered plasticity in intact animals and humans remains to be fully resolved.

Together, our work demonstrates a critical role for PRRT2 in the regulation of synaptic morphology in the 16p11.2 duplication, and provides further evidence that dysfunctional homeostatic plasticity could contribute to mental disorders.

## Acknowledgments

We would like to thank Professor Nicolas Katsanis (Duke University) for providing the backcrossed *16p11.2^dup/+^*mice. *Prrt2*^+/-^ mice were kindly provided by the University of Tennessee Health Science Center.

## Authors contributions

M.P.F. and P.P. designed research; M.P.F., N.H.P., V.A.B., L.E.D., S.Y., and M.D.S. performed research; M.P.F., N.H.P., V.A.B., L.E.D., S.Y., M.D.S. analyzed data; M.S.L. generated and provided mice, and M.P.F. and P.P. wrote the manuscript.

## Data and materials availability

Any materials generated in this study are available upon request.

## Methods

### Mouse models

All animal procedures were performed with the approval of the Institutional Animal Care and Use Committee (IACUC) at Northwestern University (IS00006149 and IS00015320). The 16p11.2^dup/+^ mouse model carrying a duplication of mouse chromosome 7qF4 (the syntenic region of human chromosome 16p11.2) was generated by Dr. Alea Mills (Cold Spring Harbor Laboratory) (47). Mice were backcrossed >5 generations to a C57BL/6J background by Dr. Nicolas Katsanis and kindly gifted to our laboratory. Male 16p11.2^dup/+^ mice were then crossed for at least 2 generations on a C57BL/6N background (Charles River Laboratories) prior to experiments. *Prrt2*^+/-^ on a C57BL/6J background were generated from *Prrt2*^fl/fl^ and kindly donated by Mark LeDoux (48). For *Prrt2* correction experiments, 16p11.2^dup/+^ mice were crossed with *Prrt2*^+/-^ mice. Heterozygous Thy1-YFP-H mice (Strain #:003782, C57BL/6J background) were purchased from Jackson Laboratory used at 3 months of age. Mice were housed according to gender on a 14-h on/10-h off light/dark cycle with temperature 70-74°F and humidity 30-70%. Up to five animals were housed per cage, with mixed-genotype littermates. All experiments were performed with littermate controls.

### Primary cortical neuron cultures

All experiments were performed on 17-23 day *in vitro* (DIV) primary mouse cortical neurons. Primary cortical neurons were prepared from dissociated sibling 16p11.2^dup/+^ and 16p11.2^+/+^ embryos at embryonic day 18.5. Brains were extracted from embryos, and cortices were isolated using a dissection microscope and stored in ice cold L-15 Medium (Thermo Fisher) supplemented with penicillin (50 U/ml) and streptomycin (50 μg/ml) (pen/strep, Gibco). The cortical tissue was digested in papain (Sigma; 20 units/ml) diluted in Dulbecco’s Modified Eagle Medium (DMEM, Corning #10013CV) containing EDTA (0.5 mM, Sigma) and DNaseI (2 units/mL, Sigma). Papain solution was activated 10 mins prior to enzymatic digestion with L-cysteine (1 mM) and incubated at 37°C for 20 min. Cortical tissue was mechanically dissociated in high glucose dissociating medium (DMEM containing 10% Fetal bovine serum (FBS, Corning), 1.4 mM GlutaMAX, and 6 g/L glucose). The supernatant containing dissociated cortical neurons was seeded on 12 well plates containing coverslips (Neuvitro Corporation) precoated overnight with 50 µg/ml poly-D-lysine (Sigma). Twelve-well plates contained 18 mm coverslips seeded at a density of 200’000 cells per well. Two hours after seeding the dissociation medium was replaced with Neurobasal feeding medium (Neurobasal medium supplemented with B-27 supplement (Gibco), 2 mM GlutaMAX (Gibco) and pen/strep). Neuronal cultures were maintained at 37°C in 5% CO_2_ and half media changes of Neurobasal feeding medium were performed twice a week.Rat cortical neurons were cultured similarly to mouse neurons as previously described (49).

### Lipofectamine transfections

Neurons in 12 well plates were transfected with 2-4 µg of plasmid and 2-4 µl of Lipofectamine 2000 (Invitrogen) according to manufacturer’s instructions. Coverslips containing neurons were placed into antibiotic-free feeding medium at 37°C for at least 30 min prior to transfections. Plasmids and Lipofectamine 2000 were diluted separately in 25 µl DMEM supplemented with HEPES (10 mM), incubated for 5min, then mixed thoroughly together, and incubated a final time for 20-30 minutes at 37°C before adding to cultured cells. Following a 4 h of incubation, neurons were transferred to antibiotic-containing feeding medium containing half conditioned and half fresh medium, and constructs were allowed to express for 2-3 days prior to experiments.

### Activity-deprivation assay

Neurons were transfected with a GFP plasmid and treated with either TTX (2 µM) or vehicle (Sodium citrate buffer 100 nM). After 48h, coverslips fixed and processed for immunostaining.

### Surface and total immunostaining

For surface immunostaining, coverslips were incubated in 200 ul of neurobasal medium containing an N-terminal mouse GluA1 antibody (Millipore MAB2263, 1:300) or a rabbit GFP antibody (Abcam ab6556, 1:1000) for 15 min at room temperature (RT), followed by three washes in PBS. Surface labeling was then followed by regular immunocytochemistry to label all proteins (total staining). For total staining, coverslips were fixed using 4% formaldehyde (Sigma) in 4% sucrose/PBS for 15 min at RT, and washed another three times in PBS. Neurons were permeabilized and blocked simultaneously in PBS containing 0.1% Triton and 5% normal goat serum (Jackson Immunoresearch) for 1 h at RT followed by incubation with primary antibodies overnight at 4°C. coverslips were washed three times in PBS and incubated with secondary antibodies at room temperature for 1 hr followed by another three washes in PBS. Coverslips with immunostained neurons were briefly washed in distilled water and mounted onto microscope slides using Prolong anti-fade reagent (Life Technologies). Primary and secondary antibodies were diluted in PBS containing 5% normal goat serum. Neurons from the same differentiation experiment were fixed and stained at the same time with identical antibody dilutions. Primary antibodies for total staining were as follows: Rabbit C-terminal GluA1 (Millipore AB1504, 1:300), Chicken GFP (Abcam ab13970, 1:10,000), Rabbit PRRT2 (Sigma HPA014447, 1:100), Guinea pig VGLUT1 (synaptic systems Cat# 135 304, 1:1000), Mouse PSD-95 (NeuroMab clone K28/43 1:1000). Secondary antibodies: Alexa 647 goat anti-mouse (1:1000), Alexa 647 goat anti-rabbit (1:1000), Alexa 647 goat anti-guinea pig (1:1000), Alexa 568 goat anti-mouse (1:1000) Alexa 568 goat anti-rabbit (1:1000), Alexa 488 goat anti-chicken (1:1000).

### Western blotting

Cortical lysates were prepared by homogenizing tissue with a Dounce homogenizer and solubilizing in cold RIPA buffer (50mM Tris pH 7.4, 150mM NaCl, 5mM EDTA, 1% Triton X-100, 0.5% sodium deoxycholate, 0.1% SDS, + protease inhibitors added fresh) for 1 h prior to SDS-PAGE. Proteins were transferred to PVDF membranes (Bio-Rad #170-4156) using a Trans-Blot® Turbo™ (Bio-Rad) for 15 min. Western blotting was performed with a GluA1 mouse antibody (Millipore MAB2263, 1:300) antibody.

### Neuronal morphology analysis

Images of GFP-transfected neurons were acquired on a Nikon C2+ confocal microscope with NIS-Elements software. All image acquisitions and data analyses were performed blind to genotypes. For analysis of dendritic complexity, images of neurons were acquired with a ×10 objective lens and imaged on multiple z-planes to capture full dendritic branches. Maximum intensity projections of each neuron created in Image J (National Institutes of Health, Bethesda, MD) were used to trace dendritic branches. The center of the soma was defined manually in Image J and the “Sholl Analysis” plugin was used to quantify the number of dendritic intersections occurring at each 10 μm intervals from the soma. For spine analysis and puncta analysis, multi-channel images were acquired using ×63 oil immersion objective lens. Each channel was acquired sequentially to prevent fluorescence bleed-through with 2x line averaging for each channel and 1.5x digital zoom. All images were acquired using identical settings for each channel to allow comparison between groups. Maximum intensity projections of each image were used for analysis. Spine analysis was performed in Image J using 50-100 μm segment of apical dendrite. The GFP channel was thresholded and a region of interest (ROI) containing all spines within the selected dendritic segment was analyzed. Spine area was quantified in Image J and spine density was counted manually.

### Image analysis

Image analysis was performed on the same 63 x images used for dendritic spine analysis (see Neuron morphology analysis). Analysis was performed on Image J on 50-100 μm segment of dendrite. Mean intensity values in either spine, dendrite or soma were measured in Image J using ROIs created from the cell fill (either GFP or tRFP). For puncta analysis, channels were manually thresholded to include visible puncta. Puncta analysis was performed on each channel using the ‘Analyze Particle’ function. Images of binarized puncta from each channel were then used to create a map of co-localized pixels used for co-localization analysis. The number of colocalized puncta was used to calculate the linear density.

### Live imaging of SEP-GluA1

All experiments were performed at DIV19-23 primary cortical neurons cultured on 18 mm coverslips by experimenters blind to genotype. Neurons were co-transfected with SEP-GluA1 and tRFP and constructs were allowed to express 2 days. Images were acquired on a Nikon A1B point scanning confocal fluorescence microscope (Nikon Instruments, Melville NY, USA) using a Nikon Plan Fluor 40X oil immersion objective and GaAsP point detector. Coverslips were transferred into an imaging chamber (Warner instrument, QR-41LP) with recording buffer (in mM: 150 NaCl, 2 CaCl2, 5 KCl, 10 HEPES pH 7.4, 30 glucose, 0.001 TTX, 0.01 strychnine, and 0.03 picrotoxin). The chamber was then placed in the microscope incubator (Tokai Hit, STXG-TIZWX-SET) and incubated at least 10 minutes before acquisition via a peristaltic pump (Gilson) injecting buffer at 1ml/min in the chamber. A first image stack (Ti Drive, 0,7 um step, 9 Steps, 2048×2048, Pixel Dwell 3.2) in the red and green channel was acquired then followed by a bleaching step (100% 488 laser, 8 loops for a total of 1.03 minutes duration) and a 6 image stack acquired every 5 minutes for 25 minutes. The chamber was injected with stimulation solution (recoding solution + 0.5 mM glycine with/out 50 μM APV) for 3 minutes at the beginning of the bleach step followed by recording buffer. Dendritic ROIs were created using the cell fill (tRFP) and used to measure fluorescence intensity of SEP-GluA1 in all images from the time series. Mean intensity values were normalized to the starting intensity (set to 100%) and imported into GraphPad for statistical analysis.

### Immunoprecipitation in HEK cells and mouse neocortex

HEK293T cells (ATCC) were cultured in DMEM containing 10% FBS and 1% pen/step. Cells were seeded onto 10 cm plates and transfected with lipofectamine 2000 using manufacturer’s instructions. 5μg of FLAG-*Prrt2* plasmid (GeneCopoeia) and 5μg of GluA1 plasmid were transfected and allowed to express for 48h prior to immunoprecipitation (IP). HEK293T cells were washed with PBS, homogenized in IP buffer (50 mM Tris, pH 7.4, 150 mM NaCl, 1% Triton X-100, with protease inhibitor cocktail) and solubilized for 1 h at 4 °C. Solubilized material was clarified by centrifugation at 15,000g for 10 min at 4 °C. Soluble proteins were then incubated with 1 μg of anti-FLAG antibody (F1804, Sigma) or 1 μg of mouse IgG (sc-2025, Santa Cruz) overnight at 4 °C, followed by a 1 h capture with 20μl protein A/G beads the following day. Beads were then washed three times with IP buffer before adding 2× Laemmli buffer (Biorad) and boiling at 95 °C for 5min. Mouse brain IPs were performed on a solubilized P2 membrane fraction from neocortex of adult mice (>6 weeks), as above. Input and IP Samples were analyzed by SDS-PAGE and western blotting.

### STED imaging

Rat neurons were transfected with GFP and immunostained with primary antibodies against PRRT2 (Sigma HPA014447, 1:100) and GluA1 (Millipore MAB2263, 1:300), and the following Abberior secondaries: STAR 460L (1:1000), STAR Orange (1:1000), STAR Red (1:1000). Images were acquired on an Abberior STEDycon and processed in Image J.

### Expansion microscopy

We transcardially perfused 3-month-old male Thy1-YFP-H heterozygous mice with 4% paraformaldehyde in PBS (pH 7.4). The brains were then placed in 30% sucrose/PBS overnight at 4°C. Subsequently, the brains were embedded in frozen tissue-cutting medium (Tissue-Tek O.C.T. Compound) and coronal sections were performed using a Cryostat (Leica) at a thickness of 50 microns. Expansion microscopy was performed on Thy1-YFP-H brain slices following the Damstra *et al* protocol for slices (50). Briefly, floating slices were permeabilized and blocked (PBS with 0.1% Triton, 5% BSA) for 4 hours and incubated at 4°C overnight in this buffer with primary antibodies: GFP chicken (1:1000), PRRT2 rabbit (1:100) and GluA1 mouse (1:200). Then, slices were washed 3 times for 30 minutes (PBS, 0.1% Triton) and incubate during 3 hours with the secondary antibodies: CF488 anti-chicken (1:500), CF568 anti-mouse (1:500) and CF640 anti-rabbit (1:500) (Biotium Inc, Fremont CA, USA) washed again and incubated in acryloyl-X SE (10ug/ml, Thermofisher) overnight. After one wash with PBS, slices were incubated in TREx solution for 20 minutes on ice and then for 1 hour at 37°C. The gels were incubated in proteinase K digestion buffer for 4 hours at 37°C, washed 3 times with Milli-Q water 2 times for 30 minutes, and the last one overnight. Expanded slice size was measured to calculate the expansion factor and conserved at 4°C until image acquisition. A piece of the gel including the somatosensory cortex was cut and transferred into a FluoroDish FD35 (World Precision instrument) and images were collected on a Nikon AXR point scanning confocal fluorescence microscope using a Nikon CFI Apo LWD Lambda S 40XC WI objective and GaAsP point detectors. 3D reconstructions were generated using the Imaris software (Oxford Instrument).

### Protein:protein interaction networks

Protein interaction networks were built in Cystoscape (v3.6.0) using data from published PRRT2 and AMPA receptor interactomes (10, 15, 51). All AMPA receptor-interacting proteins in the PRRT2 interactome were imported into Cystoscape, as well as all PRRT2-associated proteins involved in SNARE-complex formation.

### Golgi-Cox staining

Mice 8-10 weeks of age were euthanized and perfused with 4% paraformaldehyde in phosphate-buffered saline (PBS). Brains were removed and treated according to the manufacturer’s instructions in the FD Rapid GolgiStain Kit (FD Technologies, Columbia, MD, USA). The tissue was immersed in impregnation solution A/B for 24 h and solution C for 2 weeks prior to cryosectioning. Mounted sections of 120 µm thickness were imaged on a microscope (LSM 510; Carl Zeiss) using a 63x oil lens with AxioCam software from Zeiss (Oberkochen, Germany). Images were stacked and analyzed using ImageJ. Dendritic spine analysis was performed on secondary apical dendrites from pyramidal neurons in layer 2/3 of the primary somatosensory cortex.

### Statistical analysis

All statistical tests were performed with GraphPad Prism. All data was tested for normality using the D’Agostino and Pearson omnibus normality test. An unpaired t-test was used if the data was parametric and a Mann-Whitney test was used if the data was non-parametric. Bar graphs are displayed as mean ± SEM, unless otherwise specified. For Sholl analysis, a two-way repeated measures ANOVA was performed to detect an effect of genotype or genotype-distance interaction. A post-hoc Bonferroni correction was then performed to determine the statistical significance at each distance interval. For spine and puncta analyses with corrected mice, a two-way ANOVA with Tukey’s post-hoc test was performed with effect of *Prrt2* genotype and effect of 16p11.2 genotype as independent variables. Hypergeometric tests for gene set analysis were performed in R. p-values were considered significant if p< 0.05. For all statistical tests asterisks indicate: *, p<0.05; **p<0.01, ***, p<0.001.

## References

1. McCutcheon RA, Reis Marques T, Howes OD (2020): Schizophrenia-An Overview. JAMA psychiatry. 77:201–210.

2. Lord C, Elsabbagh M, Baird G, Veenstra-Vanderweele J (2018): Autism spectrum disorder. Lancet. 392:508–520.

3. De Rubeis S, He X, Goldberg AP, Poultney CS, Samocha K, Cicek AE, et al. (2014): Synaptic, transcriptional and chromatin genes disrupted in autism. Nature. 515:209–215.

4. Bourgeron T (2015): From the genetic architecture to synaptic plasticity in autism spectrum disorder. Nature Reviews Neuroscience. 16:551–563.

5. Trubetskoy V, Pardinas AF, Qi T, Panagiotaropoulou G, Awasthi S, Bigdeli TB, et al. (2022): Mapping genomic loci implicates genes and synaptic biology in schizophrenia. Nature. 604:502–508.

6. Forrest MP, Parnell E, Penzes P (2018): Dendritic structural plasticity and neuropsychiatric disease. Nature reviews Neuroscience. 19:215–234.

7. Mullins C, Fishell G, Tsien RW (2016): Unifying Views of Autism Spectrum Disorders: A Consideration of Autoregulatory Feedback Loops. Neuron. 89:1131–1156.

8. Ebert DH, Greenberg ME (2013): Activity-dependent neuronal signalling and autism spectrum disorder. Nature. 493:327–337.

9. Rein B, Yan Z (2020): 16p11.2 Copy Number Variations and Neurodevelopmental Disorders. Trends in neurosciences. 43:886–901.

10. Forrest MP, Dos Santos M, Piguel NH, Wang YZ, Hawkins NA, Bagchi VA, et al. (2023): Rescue of neuropsychiatric phenotypes in a mouse model of 16p11.2 duplication syndrome by genetic correction of an epilepsy network hub. Nature communications. 14:825.

11. Savino E, Cervigni RI, Povolo M, Stefanetti A, Ferrante D, Valente P, et al. (2020): Proline-rich transmembrane protein 2 () regulates the actin cytoskeleton during synaptogenesis. Cell Death Dis. 11.

12. Coleman J, Jouannot O, Ramakrishnan SK, Zanetti MN, Wang J, Salpietro V, et al. (2018): PRRT2 Regulates Synaptic Fusion by Directly Modulating SNARE Complex Assembly. Cell reports. 22:820–831.

13. Valente P, Castroflorio E, Rossi P, Fadda M, Sterlini B, Cervigni RI, et al. (2016): PRRT2 Is a Key Component of the Ca(2+)-Dependent Neurotransmitter Release Machinery. Cell reports. 15:117–131.

14. Bayes A, Collins MO, Croning MD, van de Lagemaat LN, Choudhary JS, Grant SG (2012): Comparative study of human and mouse postsynaptic proteomes finds high compositional conservation and abundance differences for key synaptic proteins. PloS one. 7:e46683.

15. Schwenk J, Harmel N, Brechet A, Zolles G, Berkefeld H, Muller CS, et al. (2012): High-resolution proteomics unravel architecture and molecular diversity of native AMPA receptor complexes. Neuron. 74:621–633.

16. Spruston N (2008): Pyramidal neurons: dendritic structure and synaptic integration. Nature reviews Neuroscience. 9:206–221.

17. Hering H, Sheng M (2001): Dendritic spines: structure, dynamics and regulation. Nature reviews Neuroscience. 2:880–888.

18. Turrigiano G (2012): Homeostatic Synaptic Plasticity: Local and Global Mechanisms for Stabilizing Neuronal Function. Csh Perspect Biol. 4.

19. Wefelmeyer W, Puhl CJ, Burrone J (2016): Homeostatic Plasticity of Subcellular Neuronal Structures: From Inputs to Outputs. Trends in neurosciences. 39:656–667.

20. Wondolowski J, Dickman D (2013): Emerging links between homeostatic synaptic plasticity and neurological disease. Frontiers in cellular neuroscience. 7:223.

21. Ramocki MB, Zoghbi HY (2008): Failure of neuronal homeostasis results in common neuropsychiatric phenotypes. Nature. 455:912–918.

22. Chen L, Li X, Tjia M, Thapliyal S (2022): Homeostatic plasticity and excitation-inhibition balance: The good, the bad, and the ugly. Current opinion in neurobiology. 75:102553.

23. Matsuzaki M, Ellis-Davies GC, Nemoto T, Miyashita Y, Iino M, Kasai H (2001): Dendritic spine geometry is critical for AMPA receptor expression in hippocampal CA1 pyramidal neurons. Nature neuroscience. 4:1086–1092.

24. Diering GH, Huganir RL (2018): The AMPA Receptor Code of Synaptic Plasticity. Neuron. 100:314–329.

25. Turrigiano GG, Leslie KR, Desai NS, Rutherford LC, Nelson SB (1998): Activity-dependent scaling of quantal amplitude in neocortical neurons. Nature. 391:892–896.

26. Blizinsky KD, Diaz-Castro B, Forrest MP, Schurmann B, Bach AP, Martin-de-Saavedra MD, et al. (2016): Reversal of dendritic phenotypes in 16p11.2 microduplication mouse model neurons by pharmacological targeting of a network hub. Proceedings of the National Academy of Sciences of the United States of America. 113:8520–8525.

27. Purkey AM, Dell’Acqua ML (2020): Phosphorylation-Dependent Regulation of Ca-Permeable AMPA Receptors During Hippocampal Synaptic Plasticity. Front Synaptic Neuro. 12.

28. Davis BA, Chen HY, Ye ZY, Ostlund I, Tippani M, Das D, et al. (2024): TCF4 Mutations Disrupt Synaptic Function Through Dysregulation of RIMBP2 in Patient-Derived Cortical Neurons. Biological psychiatry. 95:662–675.

29. Kopec CD, Li B, Wei W, Boehm J, Malinow R (2006): Glutamate receptor exocytosis and spine enlargement during chemically induced long-term potentiation. Journal of Neuroscience. 26:2000–2009.

30. Wu ZY, Xu XZ, Xi P (2021): Stimulated emission depletion microscopy for biological imaging in four dimensions: A review. Microsc Res Techniq. 84:1947–1958.

31. Valente P, Romei A, Fadda M, Sterlini B, Lonardoni D, Forte N, et al. (2019): Constitutive Inactivation of the PRRT2 Gene Alters Short-Term Synaptic Plasticity and Promotes Network Hyperexcitability in Hippocampal Neurons. Cerebral cortex. 29:2010–2033.

32. Valtorta F, Benfenati F, Zara F, Meldolesi J (2016): PRRT2: from Paroxysmal Disorders to Regulation of Synaptic Function. Trends in neurosciences. 39:668–679.

33. Wu D, Bacaj T, Morishita W, Goswami D, Arendt KL, Xu W, et al. (2017): Postsynaptic synaptotagmins mediate AMPA receptor exocytosis during LTP. Nature. 544:316-+.

34. Gu Y, Huganir RL (2016): Identification of the SNARE complex mediating the exocytosis of NMDA receptors. Proceedings of the National Academy of Sciences of the United States of America. 113:12280–12285.

35. Gu Y, Chiu SL, Liu B, Wu PH, Delannoy M, Lin DT, et al. (2016): Differential vesicular sorting of AMPA and GABA receptors. Proceedings of the National Academy of Sciences of the United States of America. 113:E922–E931.

36. Arendt KL, Zhang YS, Jurado S, Malenka RC, Südhof TC, Chen L (2015): Retinoic Acid and LTP Recruit Postsynaptic AMPA Receptors Using Distinct SNARE-Dependent Mechanisms. Neuron. 86:442–456.

37. Kopec CD, Real E, Kessels HW, Malinow R (2007): GluR1 links structural and functional plasticity at excitatory synapses. Journal of Neuroscience. 27:13706–13718.

38. Suresh J, Radojicic M, Pesce LL, Bhansali A, Wang J, Tryba AK, et al. (2016): Network burst activity in hippocampal neuronal cultures: the role of synaptic and intrinsic currents. Journal of neurophysiology. 115:3073–3089.

39. Penn Y, Segal M, Moses E (2016): Network synchronization in hippocampal neurons. Proceedings of the National Academy of Sciences of the United States of America. 113:3341–3346.

40. Verstraelen P, Pintelon I, Nuydens R, Cornelissen F, Meert T, Timmermans JP (2014): Pharmacological characterization of cultivated neuronal networks: relevance to synaptogenesis and synaptic connectivity. Cell Mol Neurobiol. 34:757–776.

41. Bertero A, Liska A, Pagani M, Parolisi R, Masferrer ME, Gritti M, et al. (2018): Autism-associated 16p11.2 microdeletion impairs prefrontal functional connectivity in mouse and human. Brain : a journal of neurology. 141:2055–2065.

42. Genç Ö, An JY, Fetter RD, Kulik Y, Zunino G, Sanders SJ, et al. (2020): Homeostatic plasticity fails at the intersection of autism-gene mutations and a novel class of common genetic modifiers. eLife. 9.

43. Zhang ZJ, Marro SG, Zhang YS, Arendt KL, Patzke C, Zhou B, et al. (2018): The fragile X mutation impairs homeostatic plasticity in human neurons by blocking synaptic retinoic acid signaling. Science translational medicine. 10.

44. Soden ME, Chen L (2010): Fragile X Protein FMRP Is Required for Homeostatic Plasticity and Regulation of Synaptic Strength by Retinoic Acid. Journal of Neuroscience. 30:16910–16921.

45. Ellingford RA, Panasiuk MJ, de Meritens ER, Shaunak R, Naybour L, Browne L, et al. (2021): Cell-type-specific synaptic imbalance and disrupted homeostatic plasticity in cortical circuits of ASD-associated haploinsufficient mice. Molecular psychiatry. 26:3614–3624.

46. Fink JJ, Schreiner JD, Bloom JE, James J, Baker DS, Robinson TM, et al. (2021): Hyperexcitable Phenotypes in Induced Pluripotent Stem Cell-Derived Neurons From Patients With 15q11-q13 Duplication Syndrome, a Genetic Form of Autism. Biological psychiatry. 90:756–765.

47. Horev G, Ellegood J, Lerch JP, Son YE, Muthuswamy L, Vogel H, et al. (2011): Dosage-dependent phenotypes in models of 16p11.2 lesions found in autism. Proceedings of the National Academy of Sciences of the United States of America. 108:17076–17081.

48. Calame DJ, Xiao J, Khan MM, Hollingsworth TJ, Xue Y, Person AL, et al. (2020): Presynaptic PRRT2 Deficiency Causes Cerebellar Dysfunction and Paroxysmal Kinesigenic Dyskinesia. Neuroscience. 448:272–286.

49. Smith KR, Kopeikina KJ, Fawcett-Patel JM, Leaderbrand K, Gao R, Schurmann B, et al. (2014): Psychiatric risk factor ANK3/ankyrin-G nanodomains regulate the structure and function of glutamatergic synapses. Neuron. 84:399–415.

50. Damstra HGJ, Mohar B, Eddison M, Akhmanova A, Kapitein LC, Tillberg PW (2022): Visualizing cellular and tissue ultrastructure using Ten-fold Robust Expansion Microscopy (TREx). eLife. 11.

51. Shanks NF, Savas JN, Maruo T, Cais O, Hirao A, Oe S, et al. (2012): Differences in AMPA and kainate receptor interactomes facilitate identification of AMPA receptor auxiliary subunit GSG1L. Cell reports. 1:590–598.

